# Myeloid-derived suppressor cell subsets drive glioblastoma growth in a sex-specific manner

**DOI:** 10.1101/2020.02.19.954552

**Authors:** Defne Bayik, Yadi Zhou, Chihyun Park, Changjin Hong, Daniel Vail, Daniel J. Silver, Adam J. Lauko, Gustavo A. Roversi, Dionysios C. Watson, Alice Lo, Tyler J. Alban, Mary McGraw, Mia D. Sorensen, Matthew M. Grabowski, Balint Otvos, Michael A. Vogelbaum, Craig M. Horbinski, Bjarne Winther Kristensen, Ahmad M. Khalil, Tae Hyun Hwang, Manmeet S. Ahluwalia, Feixiong Cheng, Justin D. Lathia

## Abstract

Myeloid-derived suppressor cells (MDSCs) that block anti-tumor immunity are elevated in glioblastoma (GBM) patients. However, the distinct contribution of monocytic (mMDSC) versus granulocytic (gMDSC) subsets has yet to be determined. We observed that mMDSCs were enriched in the male tumor microenvironment, while gMDSCs were elevated in the circulation of female GBM models. Depletion of peripheral gMDSCs extended the survival only in female mice. Using gene expression signatures coupled with network medicine analysis, we demonstrated in pre-clinical models that mMDSCs could be targeted with anti-proliferative agents in males, whereas gMDSC function in females could be inhibited by IL-1β blockade. Analysis of patient data confirmed that proliferating mMDSCs were the predominant population in male tumors, and that a high gMDSC/IL-1β gene signature correlated with poor prognosis of female patients. These findings demonstrate that MDSC subsets differentially drive immune suppression in a sex-specific manner and can be leveraged for therapeutic intervention in GBM.

**Statement of Significance:** Sexual dimorphism at the level of MDSC subset prevalence, localization and gene expression profile comprises a therapeutic opportunity. Our results indicate that chemotherapy can be used to target mMDSC in males, while IL-1 pathway inhibitors can provide benefit to females through blockade of gMDSC function.

## Introduction

Glioblastoma (GBM) is the most common primary malignant brain tumor with a median survival of 20 months post-diagnosis^1^. Epidemiological studies further point to a 1.6-fold higher incidence among men, suggesting a male-dominant sexual dimorphism in GBM^2^. Additionally, male patients have a worse prognosis than females, underscoring the clinical relevance of studying biological sex in GBM^3^. This sex bias in part is dictated by tumor cell-intrinsic mechanisms. In particular, genome-wide association studies (GWAS) led to identification of distinct risk variants associate with GBM in males versus female patients^2^. Moreover, differences in tumor mutational profile driving oncogenic transformation and cellular metabolism were linked to the discrepancies observed in disease manifestation, outcome and susceptibility to treatment^4-7^. However, the contribution of host factors to sexual dimorphism in GBM is yet to be investigated. Notably, immunological response significantly differs between males and females. Regulated by the combination of sex chromosomes, hormones and environmental factors, sex-based differences in immunity determine the efficacy of vaccines, susceptibility to autoimmune diseases and response to infectious agents^8^. Nevertheless, it is unclear how this immunological variation affects tumorigenesis or immunotherapy response in GBM.

Myeloid cells are a major contributor of the GBM microenvironment and comprise a therapeutic target to reverse the immunosuppression that drives tumor progression^9^. Correspondingly, myeloid-derived suppressor cells (MDSCs) are enhanced in the peripheral circulation of patients with GBM compared to those with low grade central nervous system (CNS) tumors; and increased tumor infiltration of MDSCs associates with poor GBM outcome^10-13^. Initially recognized for their ability to suppress anti-tumor immune response, MDSCs are a heterogenous population of bone marrow-derived immature myeloid cells subclassified into monocytic (mMDSC) and granulocytic (gMDSC) subsets. While both MDSC subsets hinder the activity of T and natural killer (NK) cells; mMDSCs and gMDSCs can undertake additional roles in the maintenance of primary tumors versus promotion of metastasis^14,15^. Despite growing evidence implicating the distinct roles of MDSC subsets in disease progression, little is known about their differential activity in GBM. Here, we report that MDSC subset variation is an underlying factor contributing to sexual dimorphism in GBM prognosis. Our data demonstrate that mMDSCs localize to the tumor microenvironment and support GBM progression particularly in males. In contrast, systemic gMDSC accumulation is the dominant mechanism regulating the anti-tumor immune response in females. We further show that unique characteristics of MDSC subsets determine their susceptibility to distinct drug candidates, whose therapeutic efficacy was governed by host sex.

## Results

### mMDSCs primarily localize to GBM tumors, while gMDSCs accumulate in peripheral circulation

To determine the dynamics of MDSC subset accumulation, we initiated mouse GBM tumors via intracranial injection of tumor cells (**Fig. 1A**). Analysis of the immune profile of the tumor-bearing hemisphere indicated an overall increase in CD45+ cells 14 and 21 days after tumor-implantation, compared to the contralateral hemisphere (**Fig. 1B, Supplementary Fig. 1A**). Specifically, there was an increase in the frequency of mMDSCs with a concomitant decrease in gMDSCs in the tumor microenvironment (**Fig. 1C, Supplementary Fig. 1B**), skewing the mMDSC:gMDSC ratio compared to sham controls (**Fig. 1D-E**). Based on recent studies demonstrating sex differences in GBM, we further assessed MDSC subsets in male and female tumor models and found that mMDSCs accumulated particularly in male tumors leading to a significantly higher mMDSC/gMDSC ratio compared to females (**Fig. 1F-H, Supplementary Fig. 1C-E**). We also analyzed the peripheral immune response as it has been demonstrated that systemic immunity plays a critical role in in immunotherapy response^16^. gMDSCs were the predominant population in blood, and both MDSC subsets were more abundant in non-tumor bearing male animals. However, there was a further increase in the peripheral gMDSC frequency post-tumor implantation particularly driven by the female tumor-bearing animals (**Fig. 1I-K, Supplementary Fig. 1F**). These sex differences in MDSC profile extended to survival differences as female mice experienced a greater survival compared to male mice (**Supplementary Fig. 1G-H**), which has recently been reported for patients with GBM ^3^. To evaluate whether immunological variation drives a sex difference in survival, we performed bone marrow transplantation experiments with female hosts. Reconstitution of female hosts with male donor bone marrow decreased survival compared to female to female transplant controls (**Supplementary Fig. 2**), demonstrating that the male immune system has a tumor-supportive role. To determine whether these sex differences in immune profile were limited to MDSCs, we assessed other immune cell populations, including macrophage, dendritic cells, natural killer cells and T lymphocytes. There were limited differences in other tumor-infiltrating immune cell populations between male and female hosts, none of which was conserved among the two glioma models (**Supplementary Fig. 3**). We also observed that there were differences in the abundance of systemic CD4^+^ T cells between males and females (**Supplementary Fig. 4**). Collectively, these findings point to sex differences in the regulation of anti-tumor immunity, which presents as distinct MDSC subset abundance and compartmentalization.

**Figure 1.**
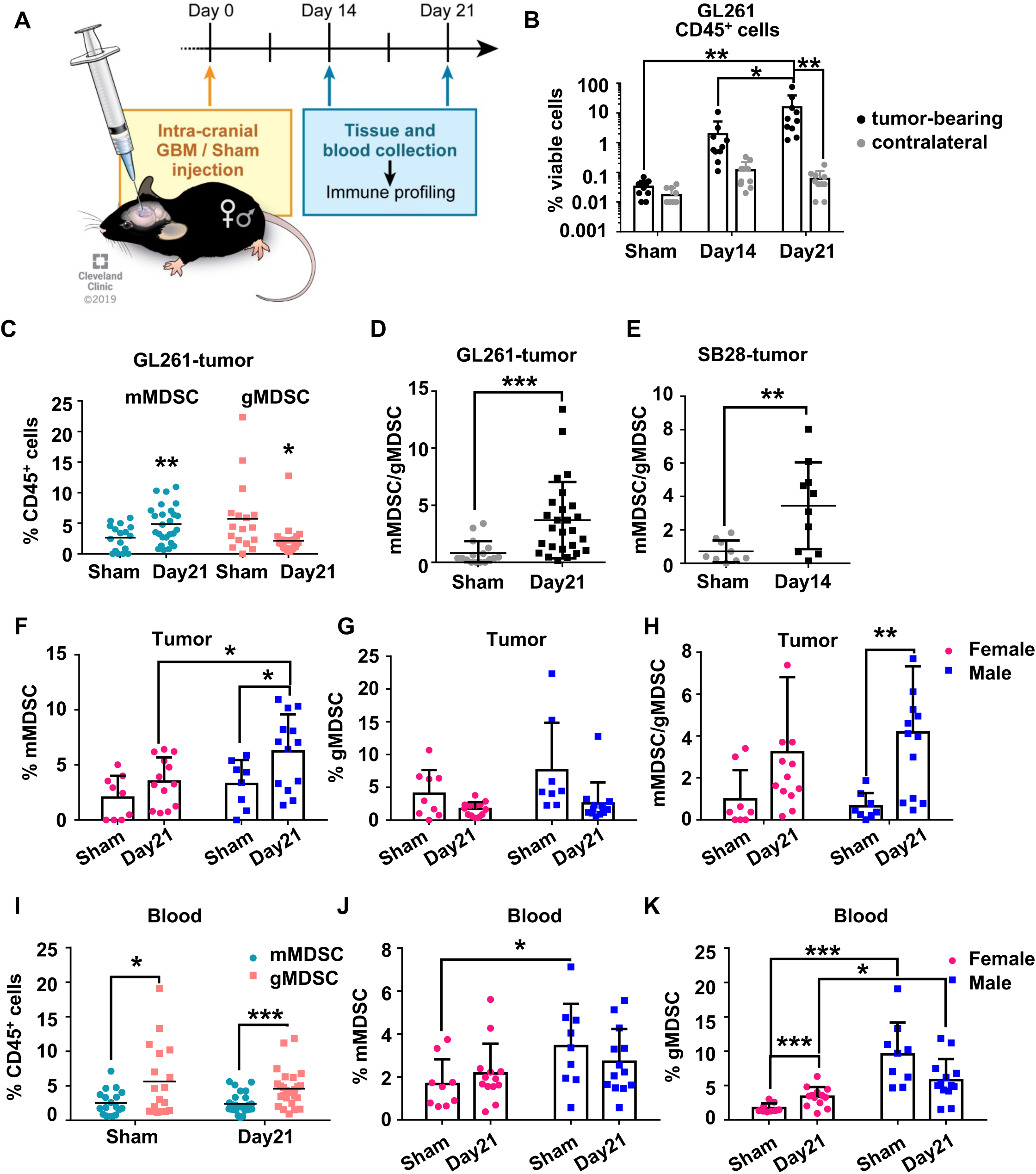
mMDSCs accumulate in tumors of male mice, while gMDSCs accumulate systemically. **A**, Timeline of immune infiltration analysis. **B**, Frequency of CD45^high^ bone marrow-derived immune cells in resected tumors versus the contralateral hemisphere of sham-injected or 25,000 GL261-implanted mice on Day 14 and Day 21. Data shown as mean ± s.d. of n = 10/group from one of independently repeated experiments. * p<0.05, ** p<0.01 as determined by two-way ANOVA. **C**, Percentage of mMDSCs (CD11b^+^CD68^-^Ly6C^+^Ly6G^-^I-A/I-E^-^) and gMDSCs (CD11b^+^CD68^-^Ly6C^-^Ly6G^+^) in CD45^+^ cells of left hemisphere from n = 10 sham-injected and n = 10 SB28-bearing animals. Data shown for individual animals and ** p<0.01 as calculated by unpaired t-test. **D**, Ratio of mMDSCs to gMDSCs in the left hemisphere from n = 16 sham-injected and n = 26 GL261-bearing animals. Data shown for individual mice combined from three independent experiments and *** p<0.001 as calculated by unpaired t-test. **E**, Ratio of mMDSC-to-gMDSC in the left hemisphere from n = 9 sham-injected and n = 10 SB28-bearing animals. Data shown for individual mice and ** p<0.01 as calculated by unpaired t-test. **F**, Percentage of mMDSCs in the CD45^+^ immune cells infiltrating the left hemisphere 21 days post-GL261 implantation or sham injection. Data shown as mean ± s.d. from three independent experiments. n = 8-9 sham-injected and n = 13 GL261-bearing mice per sex and * p<0.05 as determined by unpaired t-test. **G**, Percentage of gMDSCs in the CD45^+^ immune cells infiltrating the left hemisphere 21 days post-GL261 implantation or sham injection. Data shown as mean ± s.d. from three independent experiments. n = 8-9 sham-injected and n = 13 GL261-bearing mice per sex. **H**, Ratio of mMDSCs to gMDSCs in the left hemisphere of sham-injected or tumor-bearing mice 21 days after the procedure. Data shown as mean ± s.d. from three independent experiments. n = 8-9 sham-injected and n = 13 GL261-bearing mice per sex and ** p<0.01 as calculated by unpaired t-test. **I**, Frequency of mMDSCs and gMDSCs in the systemic circulation of sham-injected or GL261-bearing mice. Data shown for individual mice combined from three independent experiments. n = 16 sham-injected and n = 26 GL261-bearing mice, and *** p<0.001 as determined by unpaired t-test. **J**, Percentage of mMDSCs in the circulation of sham-injected and GL261-bearing mice on Day 21. Data shown as mean ± s.d. from three independent experiments. n = 9 sham-injected and n = 13 GL261-bearing mice per sex and * p<0.05 as determined by unpaired t-test. **K**, Percentage of gMDSCs in the circulation of sham-injected and GL261-bearing mice on Day 21. Data shown as mean ± s.d. from three independent experiments. n = 9 sham-injected and n = 13 GL261-bearing mice per sex and * p<0.05 as determined by unpaired t-test.

### gMDSC depletion provides survival benefit to females, while proliferative capacity of mMDSCs hinders effective depletion

Given these differences in MDSC subset compartmentalization, we sought to determine the functional contribution of MDSC subsets by utilizing neutralizing antibodies for peripheral depletion (**Fig. 2A**). Bulk MDSC depletion resulted in a survival extension, which was exclusive to females (**Fig. 2B, C**). MDSC subset specific depletion confirmed that the increase in survival of female mice was due to gMDSC depletion (**Fig. 2D-E**; for SB28, isotype and /anti-Ly6G median survival = 19.5 and /24 days in females, respectively, *data not shown*). However, there was no extension of survival with mMDSC targeting in either sex (**Fig. 2F-G**). To confirm the efficacy of this strategy, we assessed MDSC subsets in the blood and tumors, and observed that while gMDSCs could be peripherally reduced, mMDSCs remained unaltered in the tumor microenvironment (**Supplementary Fig. 5A**). A potential explanation for the lack of mMDSC reduction is that mMDSCs are proliferating. We directly assessed this by Ki-67 staining of MDSC subsets from tumor-bearing mice, and observed that mMDSCs particularly from blood highly expressed Ki-67 compared to gMDSCs independent of sex (**Fig. 2H-I; Supplementary Fig. 5B-C**). These data demonstrate a sex-specific MDSC localization in GBM models with male tumors having enhanced mMDSC accumulation in the tumor microenvironment, and females having increases gMDSCs in the peripheral circulation (**Fig. 2J**).

**Figure 2.**
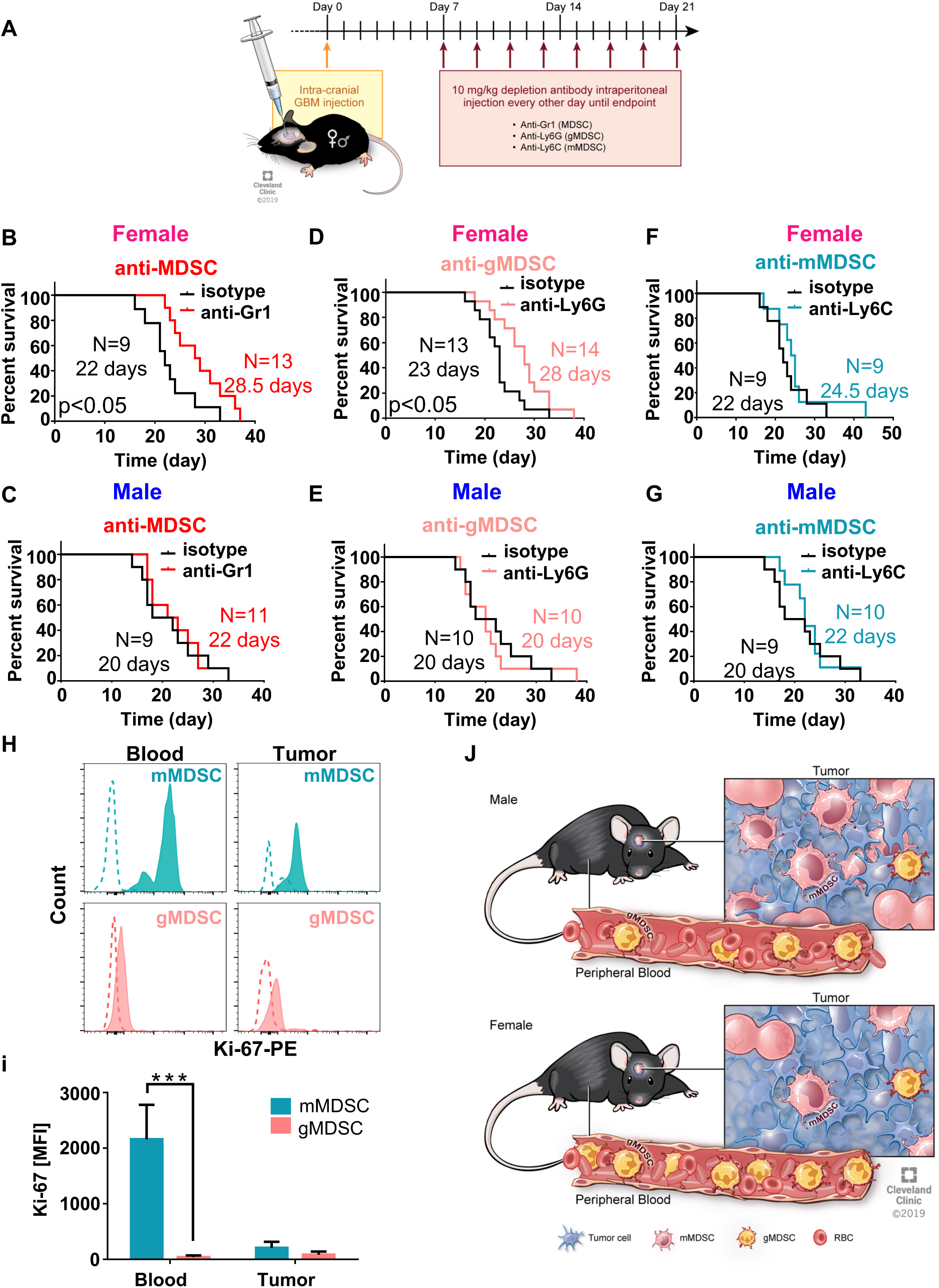
gMDSCs depletion extends the survival span of female mice. **A**, Schematics of MDSC depletion regimen. **B-G**, Kaplan-Meier curves depicting survival of female and male mice treated with (**B-C**) anti-Gr-1, (**D-E**) anti-Ly6G, and (**F-G**) anti-Ly6C neutralizing antibodies every other day starting 7 days post-GL261 implantation with respect to the isotype-treated mice. For **B**, n = 9 for isotype- and n = 13 for anti-Gr-1-treated female mice; for **C**, n = 9 for isotype- and n = 11 for anti-Gr-1-treated male mice; for **D**, n = 13 for isotype- and n = 14 for anti-Ly6G-treated female mice; for **E**, n = 10 for isotype- and n = 10 for anti-Ly6G-treated male mice; for **F**, n = 9 for isotype- and n = 9 for anti-Ly6C-treated female mice; for **G**, n = 9 for isotype- and n = 10 for anti-Ly6C-treated male mice. Data combined from two-to-three independent experiments. Significance was as determined by Gehan–Breslow–Wilcoxon test with p<0.05 being considered a significant difference. **H**, Representative histograms depicting ex vivo intracellular Ki-67 staining of mMDSCs and gMDSCs from blood and tumor. **I**, Quantification of Ki-67 staining as mean fluorescence intensity from n = 4 (2 male/female) animals euthanized on Day 21 post-GL261 implantation. Data corrected for background based on fluorescence-minus-one staining and shown as mean ± s.d. *** p<0.001 as determined by unpaired Student’s t-test. **J**, Proposed mechanism of differential mMDSC versus gMDSC accumulation in male and female mice.

### Distinct biological function of mMDSCs and gMDSCs determines their drug susceptibility

To gain mechanistic insight into the differential roles of MDSC subsets, we generated mMDSCs and gMDSCs from the bone marrow of C57BL/6 mice by adopting previously described polarization methods involving GM-CSF and IL-4Rα stimulation, which could be achieved by IL-4 or IL-13^17-19^. MDSCs generated by GM-CSF and IL-13 stimulation were functionally suppressive (**Fig. 3A-C; Supplementary Fig. 6A**). Expression profiling revealed unique gene signatures between MDSC subsets, which translated to the upregulation of distinct biological pathways (**Fig. 3D-E; Supplementary Fig. 6B-D**). While cell proliferation pathways were active in mMDSCs, consistent with increased Ki-67 expression (**Fig. 2H-I**), immune-modulatory pathways were elevated in gMDSCs. To identify putative drug targets for these subsets, we leveraged a network medicine approach that takes advantage of reported drug target interactions^20^. Using the differentially expressed gene profiles of mMDSCs and gMDSCs, we identified unique drug cohorts for each subset. Previous studies have demonstrated conflicting results of the effect of chemotherapies on MDSCs. Doxorubicin–cyclophosphamide treatment was shown to increase the frequency of MDSCs in breast cancer patients^21,22^. In contrast, nucleoside analogs were shown to reduce MDSC frequencies in various tumor types via a mechanism that was not well defined^23-26^. Our studies in mouse models, and patients with GBM, indicated that 5-fluorouracil (5-FU) and its oral formulation capecitabine could be promising candidates to target innate immunosuppression^27,28^. We focused on utilizing fludarabine for mMDSC targeting as it was predicted to be more effective than other chemotherapies (5-FU and capecitabine) previously used in GBM (**Fig. 3F, Supplementary Fig. 6E**). For gMDSC targeting, IL-1β pathway inhibitors were enriched among the top targets (**Fig. 3F, Supplementary Fig. 6F**). Consistent with the predictions of the network medicine analysis and the observation that gMDSC targeting provides survival benefit to females, we detected significantly higher IL-1β expression in gMDSC subset of female origin (**Supplementary Fig. 6G**). To test the therapeutic utility of these predicted drugs, we assessed their efficacy in pre-clinical models (**Fig. 4A**). We observed that fludarabine significantly extended survival in male mice, with no significant benefit for female mice (**Fig. 4B**). In contrast, anti-IL-1β treatment significantly prolonged the survival of female mice (**Fig. 4C**). These data support implementation of a sex-specific therapeutic strategy against GBM by targeting MDSC subsets with unique inhibitors.

**Figure 3.**
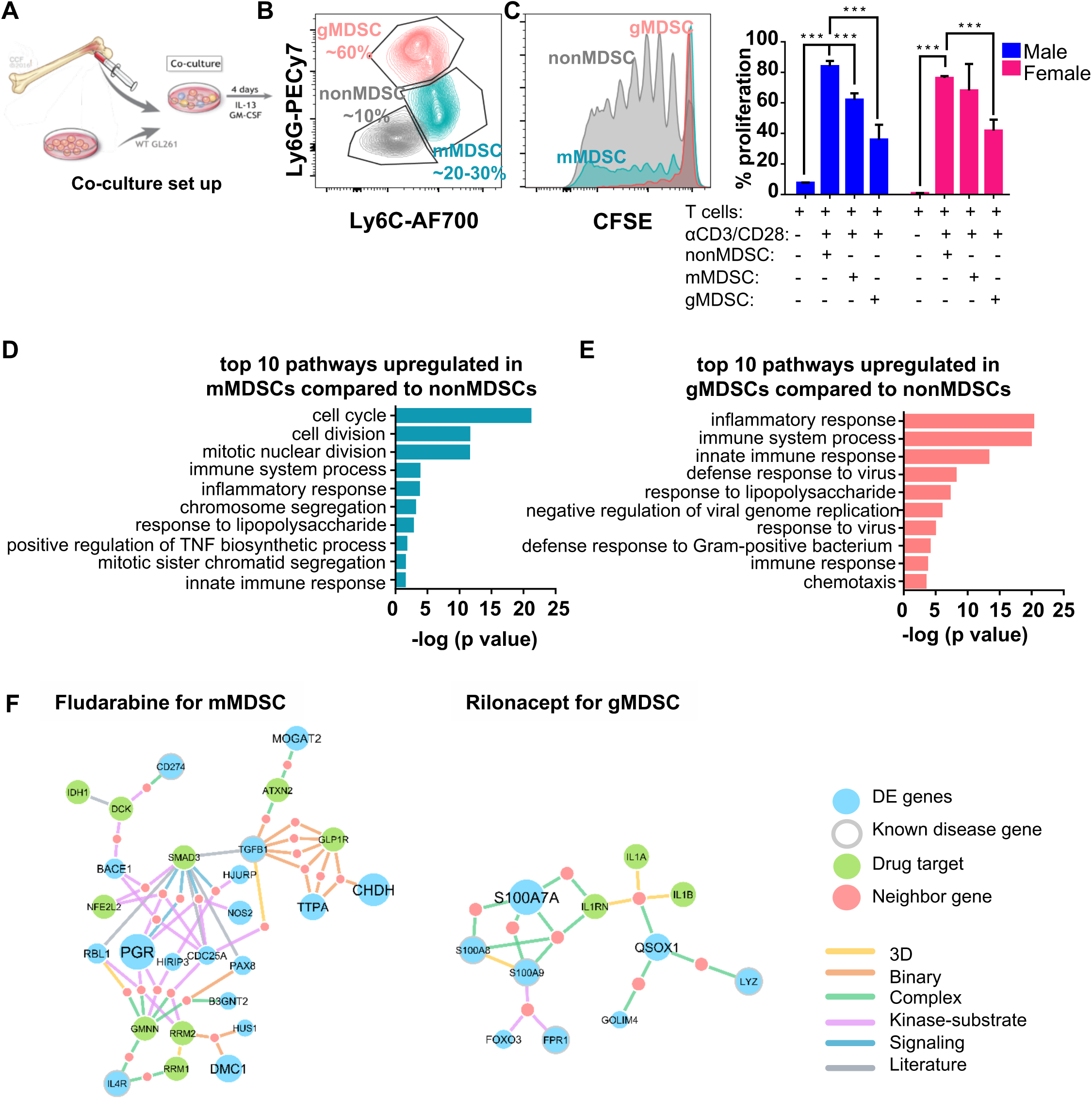
Proliferation of mMDSCs can be targeted by fludarabine, while IL-1 pathway inhibition is predicted to counteract gMDSC-mediated immunosuppression. **A**, Schematics of in vitro MDSC polarization approach. Bone marrow cells from C57BL/6 mice were co-cultured with GL261 cells in the presence of GM-CSF/IL-13 for 3 days. **B**, mMDSC, gMDSC and nonMDSC populations were phenotypically discriminated based on CD11b, Ly6C and Ly6G expression. **C**, Proliferation rates of activated T cells co-cultured with nonMDSCs, gMDSCs and mMDSCs generated by cytokine polarization, compared to unstimulated T cells. Data analyzed separately for n = 3 male and female mice and shown as mean ± s.d. *** p<0.001 as determined by two-way ANOVA. **D-E**, GeneOntology Enrichment Analysis using differentially expressed genes between (**D**) mMDSCs versus nonMDSCs and (**E**) gMDSCs versus nonMDSCs from n = 6 biological replicates based on log(fold change) <= −1 and adjusted p-value < 0.001. **F**, Potential mechanism-of-action of fludarabine and rilonacept by network inference. Differentially expressed genes of mMDSCs for fludarabine and gMDSCs for Rilonacept were directly connected to the drug targets, and/or through one or more common neighbors. For Fludarabine, a maximum distance of 2 between drug targets and differentially expressed genes was used to visualize the network. For rilonacept, the distance was set to 4. Node size indicates log_2_FC and interaction types were color coded.

**Figure 4.**
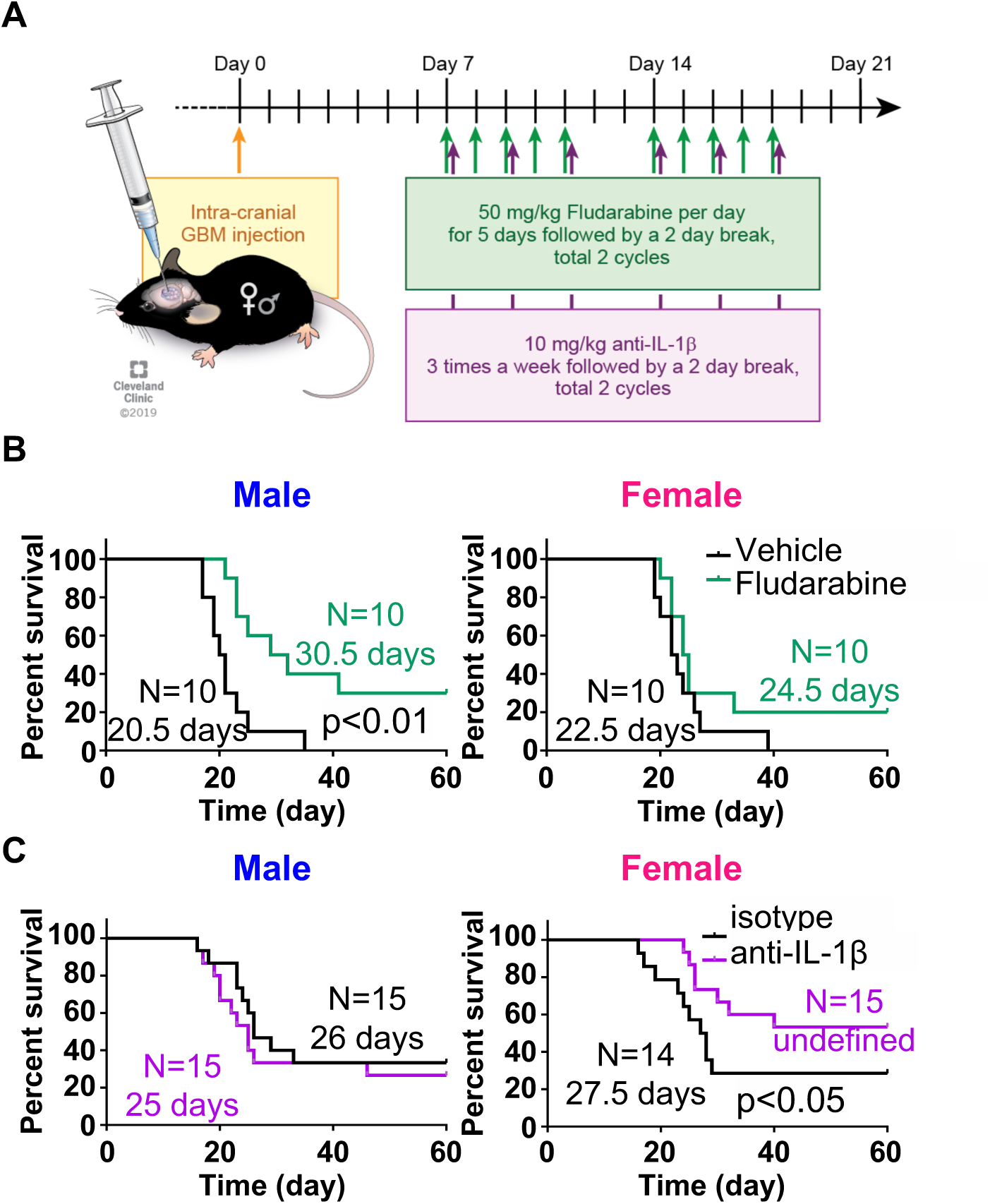
mMDSC targeting by fludarabine benefits males, while gMDSC interference with anti-IL-1β antibody improves the survival of female mice. **A**, Treatment regimen for testing the predicted drug candidates fludarabine and anti-IL-1β neutralizing antibody. **B**, Kaplan-Meier curves depicting survival of male (left) and female (right) mice treated with fludarabine. Data presented from two independent experiments with n = 10 vehicle-treated males, n = 10 fludarabine-treated males, n = 10 vehicle-treated females and n = 10 fludarabine-treated females. p<0.01 as determined by Gehan–Breslow–Wilcoxon test. **C**, Kaplan-Meier curves depicting survival of male (left) and female (right) mice treated with anti-IL1β neutralizing or isotype control antibody. Data presented from three independent experiments with n = 15 isotype control-treated males, n = 15 anti-IL-1β antibody-treated males, n = 15 isotype control-treated females and n = 15 anti-IL-1β antibody-treated females. p<0.05 as determined by Gehan–Breslow–Wilcoxon test.

### Male patients have an enhancement in tumor-infiltrating mMDSCs, while gMDSC increase associates with poor prognosis in female patients

To validate these observations in patients, we re-analyzed the immune-inhibitory myeloid cells in 188 GBM specimens in a sex-specific manner. Male patients had more IBA1+ and CD204+ cells in their tumors based on the immunohistochemistry staining (**Fig. 5A**), suggesting an increase in the immunosuppressive myeloid cell population^29^. Analysis of a separate cohort of male patients revealed that mMDSCs were the prevalent subset in human GBM tissue (**Fig. 5B-D; Supplementary Fig. 7A**). Moreover, mMDSCs were positive for Ki-67, confirming the observations that these cells are actively proliferating (**Fig. 5E-F**). To evaluate the prognostic value of gMDSCs, we analyzed The Cancer Genome Atlas (TCGA) for mRNA levels of OLR1 (LOX-1), a gMDSC-specific marker^30^, and found that high OLR1 expression inversely correlated with the survival of female but not male patients with GBM (**Fig. 6A; Supplementary Fig. 7B**). Consistently, OLR1 expression positively correlated with IL-1β expression, and high IL-1β mRNA portended poor prognosis only in female patients despite lack of sex differences in the tissue expression level of IL-1β at RNA or protein level (**Fig. 6B-E; Supplementary Fig. 7C**). Collectively, these studies indicated that proliferating mMDSCs, which are the prevalent subset in male patient with GBM, can be intervened with chemotherapies, while IL-1β represents a therapeutic target for female patients (**Fig. 6F**).

**Figure 5.**
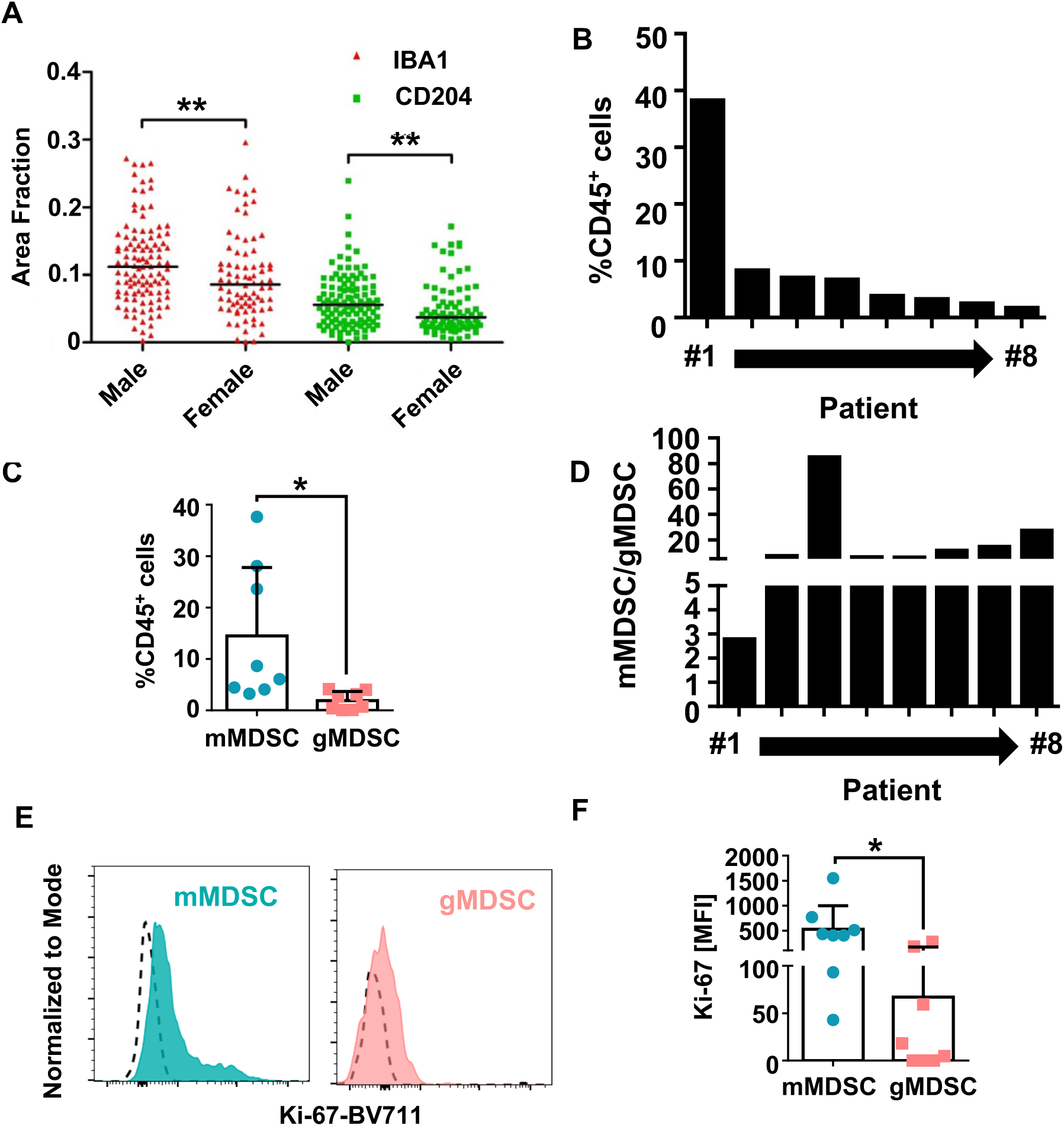
Male GBM tumors are infiltrated by proliferating mMDSCs. **A**, Fraction of IBA1-positive and CD204-positive area of n = 240 patients with primary glioma from Sorensen et al., *Neuropathology and Applied Neurobiology*, 2018, was re-analyzed by accounting for biological sex. Data shown from n = 108 males and n = 80 females as ther median. ** p<0.01 as determined by unpaired t-test. **B**, Percentage of CD45^high^ cells in viable single cells isolated from GBM tumors. Data shown for n = 8 individual patients. **C**, The frequency of mMDSCs (CD11b^+^CD33^+^CD14^+^HLA-DR^-^CD68^-^) and gMDSCs (CD11b^+^CD33^+^CD66b^+^LOX-1^+^) in tumor-infiltrating leukocytes from n = 8 male patients with GBM. Data shown as mean ± s.d. ** p<0.05 as determined by unpaired Student’s t-test. **D**, Ratio of mMDSCs to gMDSCs in tumors of n = 8 patients with GBM. **E**, Representative histograms showing Ki-67 expression levels in matched mMDSCs and gMDSCs from the same patient, in comparison to the isotype control staining. **F**, Mean fluorescence intensity of Ki-67 in tumor-infiltrating mMDSC and gMDSC from n = 8 patients with GBM. Data shown as mean ± s.d. * p<0.05 as determined by unpaired Student’s t-test.

**Figure 6.**
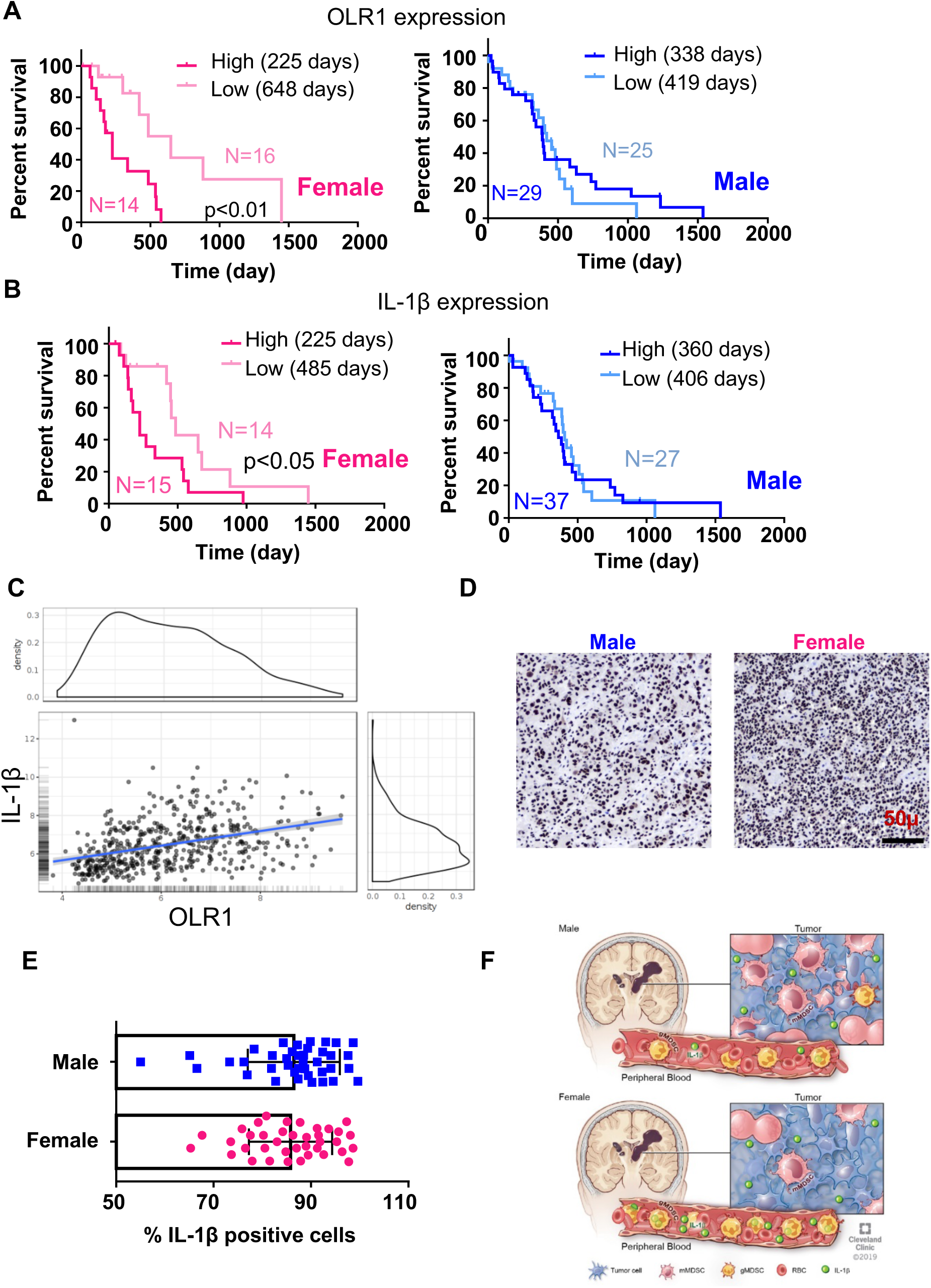
gMDSC prevalence predicts poor prognosis of female patients with GBM. **A**, Correlation among OLR1 expression levels, patient sex and survival duration was analyzed via TCGA GBM dataset. High and low expression levels were determined based on quartiles. Data shown for N = 14 female patients with high OLR1 expression, n = 16 female patients with low OLR1 expression (top) and for n = 29 male patients with high OLR1 expression, n = 25 male patients with low OLR1 expression (bottom). p<0.01 as determined by Gehan–Breslow– Wilcoxon test. **B**, Correlation among IL-1β expression levels, patient sex and survival duration was analyzed via TCGA GBM dataset. High and low expression levels were determined based on quartiles. Data shown for n = 15 female patients with high IL-1β expression, n = 14 female patients with low IL-1β expression (top) and for n = 37 male patients with high IL-1β expression, n = 27 male patients with low IL-1β expression (bottom). p<0.05 as determined by Gehan–Breslow–Wilcoxon test. **C**, Correlation between OLR1 and IL-1β expression levels from n = 538 patients. p<0.01 based on two-sided t-test. **D**, Representative images from male and female patients confirming IL-1β protein expression in GBM by immunohistochemistry. **E**, Quantification of IL-1β-positive nuclei in tumors from four visual field of n = 10 male and n = 10 female patients. **F**, Proposed model of relative mMDSC abundance and IL-1β presence in patients with GBM.

## Discussion

Immunotherapy options remain limited for GBM, despite the success of these treatment modalities in pre-clinical models and other solid tumors^31-34^. The immunosuppressive tumor microenvironment mainly consisting of myeloid cells is one of the major factors limiting the efficacy of existing treatment options^9^. Thus, understanding the variations in myeloid cell-driven immunosuppression and the associated molecular mechanisms can provide insight into the sexual dimorphism in GBM outcome. Our results demonstrated that MDSC subset heterogeneity could be leveraged for improved immunotherapy response in a sex-specific manner, and highlight the importance of assessment of sex as a biological variable in pre-clinical and clinical immunotherapy studies.

In addition to their suppressive function, MDSC subsets can execute distinct roles during the course of tumorigenesis. Ouzounova at al. elegantly demonstrated in breast cancer models that mMDSCs localize to the primary tumor, where they support cancer stem cells (CSCs); while gMDSCs promote metastatic spread to distant sites^14^. Correspondingly, our group previously reported that MDSC recruitment in GBM is driven by CSCs via the secretion of macrophage-migration inhibitory factor (MIF), suggesting a possible cross-talk between these two cell populations in multiple cancers^27^. However, this study was conducted only in females and did not distinguish between the MDSC subsets. Consistent with these earlier reports, we observed that mMDSCs was the dominant subset localizing to the GBM tissue both in mouse models and patients. Importantly, Chang and colleagues suggested that CCL2 can serve as an additional chemoattract and that it preferentially recruits CCR2+ mMDSC to the tumor microenvironment^35^. These studies provide a rationale for future assessment of key chemoattractant for differential MDSCs recruitment. Interestingly, the frequency of tumor-infiltrating mMDSCs was significantly higher in males compared to females. We previously reported that MDSC accumulation between the initial diagnosis and recurrence predicts poor GBM outcome in patients^11^. In line with this observation, mMDSC levels in pre-clinical models associated with survival duration, and male mice succumbed to disease earlier than female mice. Collectively, our observations suggested that males have a more immunosuppressive tumor microenvironment, reinforced by mMDSCs.

In contrast to the tissue-dominant localization of mMDSC, we observed that gMDSCs poorly infiltrated the tumors in pre-clinical models and patients with GBM. It is noteworthy that gMDSCs constitute the main subpopulation elevated in the circulating blood of patients with GBM ^10,12,13^ and in murine models these cells remained mainly in the periphery as well. Interestingly, males had overall higher levels of both mMDSCs and gMDSCs at baseline compared to females. Higher MDSC frequency was previous linked to the resistance of male (NZB × NZW)F1 mice to lupus^36^. Combined with our observations, these findings collectively suggest that the differences in systemic MDSC abundance could provide a mechanistic explanation to increased immunosuppression in males, in supportive of the epidemiological evidence indicating that malignancies are more frequently observed in males as opposed to inflammatory diseases, which are prevalent in females^8^. Our bone marrow transplantation studies suggest that sex differences in MDSC subset prevalence and localization have a cell-intrinsic component. Future studies will focus on revealing the mechanisms though which these sex difference emerge. While MDSC levels remained similar in between the sham-injected and tumor-bearing male mice, there was a significant increase in the systemic gMDSCs in tumor-bearing female host. Most importantly, this elevation was accompanied by reduction in the frequency of CD8^+^ T cells, pointing to systemic regulation of anti-tumor immune response in these animals. Despite significant differences in peripheral versus tumor-infiltrating T cell activation status/clonality in GBM patients treated with checkpoint inhibitors, the make-up of peripheral T cells was shown to coordinate tissue immunity in patients responding to immunotherapies^16,37,38^. Thus, our studies suggest that targeting gMDSCs in females could contribute to heightened immunotherapy response via modulation of peripheral T cells.

We also observed MDSC subsets had unique gene expression signatures and biologically active pathways that could account for the differences in their frequency and compartmentalization. Importantly, along with the modulation of immune response, the top pathways up-regulated in mMDSCs were related to cell cycle regulation. Although chemotherapies (5-FU, capecitabine and gemcitabine) were used to deplete MDSCs in pre-clinical models and patients with various cancer, including GBM; the mechanism by which these drugs has shown efficiency remained unknown^25,27,28^. Here, we demonstrate mMDSCs are actively proliferating in vivo and propose that the inhibition of cellular proliferation emerges as a likely explanation of the observed effect and mMDSC specificity of chemotherapies^23^. In contrast, gMDSCs were predicted to be more effective modulators of immune response based on the pathway analysis, as confirmed by their increased ability to suppress T cell proliferation. We leveraged these distinct gene expression signatures associate with suppression of the anti-tumor immune response to identify drug candidates that could be repurposed for cancer immunotherapy. Although, IL-1 is a prototypical pro-inflammatory cytokine, recent studies suggest that IL-1β can drive tumorigenesis by promoting a CSC phenotype, modulating the immune response and inducing tumorigenic edema^39-42^. Importantly, randomized clinical testing of canakinumab, an anti-IL-1β antibody, pointed out to a reduction in lung cancer incidence of patients with atherosclerosis^43^. We observed that IL-1β significantly expressed in the gMDSC fraction over mMDSCs, especially in females. Consistently, blockade of IL-1β significantly extended the survival of female mice, which have a gMDSC-dominant phenotype pointing to a tumor-promoting role of gMDSCs is driven by IL-1β.

In summary, our findings identify a differential MDSC subset signature and candidate mechanisms that could comprise a therapeutic opportunity for improved immunotherapy of GBM by accounting for patient sex. These results implicate that blockade of mMDSC proliferation with chemotherapies can reprogram the immunosuppressive tumor microenvironment in males. Finally, our study unveils the role of gMDSC-IL-1β axis in systemic GBM immunity, and further provide the rationale for clinical testing of IL-1β inhibitors in patients with GBM.

## Materials and Methods

### Reagents

Fluorophore-conjugated anti-Ly6C (Clone HK1.4, Catalog # 128024), anti-Ly6G (Clone 1A8, Catalog # 127618 or from BD Biosciences Catalog # 551460), anti-CD11b (Clone M1/70, Catalog # 101212), anti-CD68 (Clone FA-11, Catalog # 137024), anti-I-A/I-E (Clone M5/114.15.2, Catalog # 107606), anti-CD11c (Clone N418, Catalog # 117330), anti-CD3 (Clone 145-2C11, Catalog # 100330), anti-CD4 (Clone GK1.5, Catalog # 100422), anti-CD8 (Clone 53-6.7, Catalog # 100712), anti-NK1.1 (Clone PK136, Catalog # 108741), anti-F4/80 (Clone BM8, Catalog # 123118), anti-Ki-67 (Clone 16A8, Catalog # 652404), anti-CD45 (Clone 30-F11, Catalog # 103132), anti-CD45.1 (Clone A20, Catalog # 110728), and anti-CD45.2 (Clone 104, Catalog # 109808) antibodies were obtained from Biolegend (San Diego, CA) for analysis of mouse immune profiles.

*InVivo*MAb anti-mouse Ly6C (clone Monts1, catalog # BE0203), *InVivo*MAb anti-mouse Ly6G (clone 1A8, catalog # BE0075-1), *InVivo*MAb anti-mouse Gr-1 (clone RB6-8C5, catalog # BE0075), *InVivo*MAb anti-mouse/rat IL-1β (clone B122, catalog # BE0246), *InVivo*MAb rat IgG2a, anti-trinitrophenol (clone 2A3, catalog # BE0089), *InVivo*MAb rat IgGb, anti-keyhole limpet hemocyanin (clone LTF-2, catalog # BE0090), and *InVivo*MAb polyclonal Armenian hamster IgG (catalog # BE0091) were purchased from BioXCell (West Lebanon, NH) for depletion/targeting of MDSCs.

Fludarabine (Sagent Pharmaceuticals, Schaumburg, IL), sulfatrim (Pharmaceutical Associates In., Greenville, SC), isoflurane (Piramal Critical Care Inc., Bethlehem, PA), xylazine (Akorn Inc., Lake Forest, IL) and ketamine (Zoetis Inc., Kalamazoo, MI) were obtained from the Department of Pharmacy, Cleveland Clinic.

For analysis of human GBM immune profile, anti-CD3 (Clone SP34-2, Catalog # 557757, BD Biosciences), anti-CD11b (Clone ICRF44, Catalog # 557743, BD Biosciences), anti-CD33 (Clone WM53, Catalog # 562492, BD Biosciences), anti-CD66b (Clone G10F5, Catalog # 561650, BD Biosciences), anti-CD14 (Clone M5E2, Catalog # 558121, BD Biosciences), anti-HLA-DR (Clone L243, Catalog # 307638, Biolegend), anti-CD68 (Clone Y1/82A, Catalog # 333814, Biolegend), anti-LOX-1 (Clone 15C4, Catalog # 358606, Biolegend), anti-CD45 (Clone HI30, Catalog # 560777, BD Biosciences), anti-Ki67 (Clone B56, Catalog # 563755, BD Biosciences) and BV711 Mouse IgG1, k isotype control (Clone X40, Catalog # 563044, BD Biosciences) antibodies were used.

### Cell lines

The GL261 cell line was obtained from the Developmental Therapeutics Program, National Cancer Institute. SB28 cells were gifted by Dr. Hideho Okada (University of California San Francisco). All cell lines were treated with 1:100 MycoRemoval Agent (MP Biomedicals, Solon, OH) upon thawing, and routinely tested for *Mycoplasma spp*. (Lonza, Walkersville, MD). Cells were maintained in RPMI 1640 (Media Preparation Core, Cleveland Clinic) supplemented with 10% FBS (ThermoFisher Scientific, Walthman, MA) and 1 % Pen/Strep (Media Preparation Core). Cells were not grown for more than 10 passages.

### Mice

All experiments were approved by the Cleveland Clinic Institutional Animal Care and Use Committee and performed in accordance with the established guidelines. Four-weeks-old C57BL/6 mice (JAX Stock #000664) or B6 CD45.1 (B6.SJL-Ptprc^a^ Pepc^b^/BoyJ; JAX Stock #002014) were purchased from the Jackson Laboratory as required and housed in the Cleveland Clinic Biological Research Unit Facility under a 12:12 light/dark cycle.

Four- to eight-week-old C57BL/6 mice intracranially injected with 10,000-25,000 GL261 cells, 40,000 CT-2A cells, or 20,000 SB28 cells in 5 μl RPMI null media into the left hemisphere 2 mm caudal to the coronal suture, 3 mm lateral to the sagittal suture at a 90° angle with the murine skull to a depth of 2.5 mm. A portion of age- and sex-matched animals were injected with 5 μl RPMI null media to be used as sham controls. Mice were monitored daily for neurological symptoms, lethargy and hunched posture that would qualify as sign of tumor burden.

Seven days post-tumor implantation, mice were randomly assigned to control or treatment groups. For the depletion studies, mice were intraperitoneally injected with 10mg/kg neutralizing antibodies or isotype control in 100 μl PBS every other day until experimental endpoint, not more than 20 times. For targeting of IL-1β, mice were intraperitoneally injected with 10mg/kg targeting antibody or isotype control three times a week for two cycles. Fludarabine (50 mg/kg) was administered intraperitoneally daily with a regimen of 5 days treatment and 2 days break for two consecutive cycles.

### Murine tissue harvest

At pre-defined time points or humane endpoints, tumor-bearing or sham-injected mice were euthanized. Blood was collected directly into EDTA-coated Safe-T-Fill® micro capillary blood collection tubes (RAM Scientific, Nashville, TN) via cardiac puncture. Tubes were centrifuged at 1000 *g* for 10 minutes to isolate serum, and cell pellets were stained for flow. Femur and tibia were flushed with 10ml of PBS for isolation of bone marrow cells. Single-cell suspensions were prepared from organs by mincing in 10% RPMI, strained on a 40µm filter (FisherBrand), washed with PBS, counted and stained for flow cytometry analysis.

### Flow cytometry

Harvested tissue and blood were transferred into 96-well round-bottom plates (ThermoFisher Scientific) and washed twice with 200 µl PBS twice. Samples were stained with LIVE/DEAD Fixable Stains (ThermoFisher Scientific) diluted 1:1000 for 10 minutes on ice. Following a wash step, cells were resuspended in FcR Blocking Reagent (Miltenyi Biotec) at a 1:25 dilution in PBS/2% BSA (Sigma-Aldrich, St. Louis, MO) for 10 minutes on ice. Fluorophore-conjugated antibodies diluted 1:50 were added at a 1:2 ratio, and cells were further incubated for 20 minutes on ice. Samples were washed with PBS/BSA and fixed overnight in eBioscience™ Foxp3/Transcription Factor Fixation Buffer. Samples were acquired with a BD LSR Fortessa (BD Biosciences), and FlowJo (Version 10.5.0, FlowJo LLC, Ashland, OR) was used for analysis of the staining data.

### Bone marrow transplantation

Four weeks old C57BL/6 female mice were subjected to whole-body irradiation. Radiation (12 Gy) was given in two fractions with 3-4 hours apart. Reconstitution was achieved by retro-orbital injection of 2 x10^6^ bone marrow cells from B6 Cd45.1 mice. Drinking water was supplemented with Sulfatrim during the first 10 days, and mice were monitored for an additional 3 weeks for weight loss and infection symptoms before tumor implantation. Survival analysis was performed as described above.

### MDSC co-culture

Bone marrow was isolated from the femur and tibia of 8-12-week-old mice. 2 million bone marrow cells were co-cultured with 1 × 10^5^ GL261 cells in 6-well plates in 2 ml RPMI/10% FBS supplemented with 40 ng/ml GM-CSF and 80 ng/ml IL-13 (PeproTech, Rocky Hill, NJ) for 3-4 days. For validation experiments, bone marrow cells were incubated with the cytokine combination alone. Cells were stained for viability, blocked with Fc receptor inhibitor and stained with a combination of CD11b, Ly6C and Ly6G for sorting of MDSC subsets (mMDSCs: CD11b^+^Ly6C^+^Ly6G^-^ versus gMDSCs: CD11b^+^Ly6C^-^Ly6G^+^) and control (CD11b^+^Ly6C^-^Ly6G^-^) population using a BD FACSAria II (BD Biosciences).

### T cell proliferation assay

T cells were isolated from splenocytes of 8-12-week-old mice using a Pan T Cell Isolation Kit II (Miltenyi Biotec) with >90% purity. T cells were stained with 1 µM carboxyfluorescein succinimidyl ester (CFSE, Biolegend) for 5 minutes at 37°C and washed with ice cold 10% RPMI twice. A total of 100.000 T cells were co-cultured with 50,000 sorted myeloid cells in the presence of 30IU recombinant hIL-2 plus anti-CD3/CD28 Dynabeads (ThermoFisher Scientific) for 3-4 days in round-bottom plates. Samples were stained with anti-CD11b and anti-CD3. CFSE dilution was analyzed from CD3^+^CD11b^-^ cells using a BD LSR Fortessa.

### RNA Sequencing

RNA was isolated from a minimum of 2 × 10^6^ sorted MDSCs or nonMDSCs using an RNAeasy Mini Kit (Qiagen, Germantown, MD). RNA sequencing was performed by GENEWIZ (South Palinfield, NJ) using Illumina HiSeq, 2×150 bp configuration and ≥350M raw paired-end reads. An average 40.8M paired-end reads are sequenced across 18 samples. After Illumina universal adapters were trimmed, the average read length was shortened to 136 bp and 40.6M reads were kept for the downstream analysis. The RNA-seq reads were converted to transcriptome abundant matrix in the format of TPM (Transcripts Per Kilobase Million)^44^ via kallisto v0.44.0^45^ in default parameters. The kallisto reference genome sequence index was built on a mouse reference transcript sequences (GRCm38.p5) with the gene model annotation, gencode.vM16.annotation.gtf. The differential gene expression analysis was conducted with DESeq2 v1.18.1^46^ in a default setting as suggested. Then, we selected genes of which the absolute log2 fold change value is higher than or equal to 1 and Benjamini and Hochberg FDR (false discovery rate) < 0.001 and queried them into DAVID website for a functional enrichment analysis^47^. Both the biological process in the Gene Ontology and KEGG pathway (P-value <0.001) are used for functional annotation.

### Network Medicine Analysis

Gene sets for network analysis were generated by determining the overlapping differentially expressed genes between MDSC subsets and for each MDSC subset, in comparison to the nonMDSC fraction from 3 samples. Reads were quantified using Salmon (v0.9.1)^48^ quasimapping with an index created from ENSEMBL reference non-coding RNA and cDNA transcriptomes (GRCm38 release 95). Differential expression was calculated using DESeq2 (v1.20.0)^46^ after collapsing transcript level quantifications to gene level quantifications. Secondary elimination was done for mMDSCs based on read count >100 and for gMDSCs using the top 1500 highly expressed genes based on average read count. The full gene list is provided in *Supplementary Table 1*. Lineage markers, macrophage and dendritic cell markers were used as negative controls. Positive controls were determined based on previously published profiles^14,15^.

The network proximity of the drug targets and differentially expressed genes was computed using the closest method based on the human interactome:

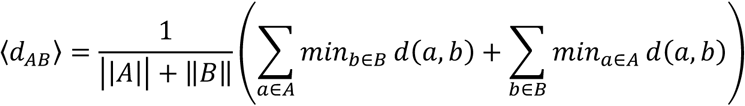

where *d*(*a, b*) is the shortest path between gene *a* and *b* from gene list *A* and *B*, respectively. The proximity was then converted to Z-score using the following method:

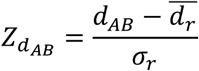

where 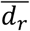 and *σ*_*r*_ are the mean and standard deviation of a permutation test repeated 1,000 times, each time using two randomly selected gene lists that have similar degree distributions to the DE genes and drug targets. A cutoff of −1 was used. Drugs were ranked based on their Z-scores. Networks were visualized using Cytoscape v3.7.1 (https://cytoscape.org/).

### Quantitative reverse-transcriptase PCR

RNA from nonMDSC, mMDSC and gMDSC fraction was isolated using RNeasy mini kit, and cDNA was synthesized with qSCRIPT cDNA Super-mix (Quanta Biosciences). qPCR reactions were performed using an Applied Biosystems™ StepOnePlus™ Real-Time PCR System and Fast SYBR-Green Mastermix (ThermoFisher Scientific). For qPCR analysis, the threshold cycle (CT) values for IL-1β was normalized to expression levels of CycloA by nonMDSCs. The following primers (Integrated DNA Technologies) were used:

IL-1β:

Forward 5’-ACGGACCCCAAAAGATGAAG-3’

Reverse 5’-TTCTCCACAGCCACAATGAG-3’

CycloA:

Forward 5’-GCGGCAGGTCCATCTACG-3’

Reverse 5’-GCCATCCAGCCATTCAGTC-3’

### Immunofluorescence

Specimens from 188 patients with a primary glioma diagnosis were stained with ionized calcium-binding adaptor molecule-1 (IBA-1) and cluster of differentiation 204 (CD204)^29^. The published dataset was reanalyzed by accounting for patient sex, and the IBA-1 and CD204 staining intensities were graphed separately for male versus female patients.

### Flow analysis of GBM tumors

All the specimens were collected by the Rose Ella Burkhardt Brain Tumor and Neuro-Oncology Center in accordance with the Institutional Review Board (IRB2559) of Cleveland Clinic. Patient demographic information is provided in *Supplementary Table* 2. Tumors were cut into small pieces with a razor blade and incubated with collagenase IV (STEMCELL Technologies, Kent, WA) on a rotator at 37°C for 1 hour. Cells were strained over a 40-µm filter and further minced with a plunger to obtain single cell suspensions. Samples were washed with 30 ml PBS twice and treated with RBC Lysis Buffer (Biolegend). Samples were stained with LIVE/DEAD Fixable Stains for 10 minutes on ice and incubated with FcR Blocking Reagent for 15 minutes on ice. Staining with fluorophore-conjugated antibodies was performed in Brilliant Stain Buffer (BD Biosciences) for 20 minutes on ice. Cells were fixed overnight in eBioscience™ Foxp3/Transcription Factor Fixation Buffer. Isotype and Ki-67 staining was performed in eBioscience™ Foxp3/Transcription Factor Permeabilization Buffer with 20 minutes of incubation at room temperature. Samples were acquired with a BD LSR Fortessa.

### Immunohistochemistry

All the samples were collected and processed at the Northwestern University Feinberg School of Medicine in accordance with approved IRB guidelines. Specimens from 10 male and 10 female patients diagnosed with GBM were stained with 1:250 diluted anti-IL-1β antibody (Clone ab156791, Abcam, Cambridge, MA). Patient demographic information and molecular characteristics of the tumors are provided in *Supplementary Table 3*.

To quantify IL-1β staining, slides were scanned with a Leica SCN400 (Leica, Buffalo Grove, IL) at 40x magnification. Four random 5000×5000 micron sections were extracted from each tissue with Aperio ImageScope (Leica). The number of IL-1β-positive cells per visual field was counted with Fiji-ImageJ software and normalized to the total number of cells (https://imagej.net/Fiji).

### The Cancer Genome Atlas

The Cancer Genome Atlas (TCGA) GBM dataset was accessed via https://xenabrowser.net/heatmap/ on 6/24/2019 and 8/5/2019 for extraction of patient sex, overall survival, IL-1β and OLR1 (LOX-1) expression level information. Survival duration was graphed for patients with highest (1^st^ quartile) versus lowest (4^th^ quartile) IL-1β and OLR1 expression. Correlation between IL-1β and OLR1 expression was determined using the https://www.showmeshiny.com/gliovis/ database.

## Statistical analysis

GraphPad PRISM (Version 6, GraphPad Software Inc., San Diego, CA) software was used for data presentation and statistical analysis. Unpaired t-test, paired t-test, two-way ANOVA were used for comparison of differences among sample groups. The Gehan–Breslow–Wilcoxon test was used to analyze survival data. The specific statistical method employed for individual data sets is listed in the figure legends.

## Conflict Statement

The authors declare no conflict of interest.

## Ethics statement

All studies performed with animals and human specimens are in compliance with the ethical regulations and approved by the respective institutional compliance offices.

## Acknowledgement

The authors thank the members of the Lathia Lab for insightful discussions. We would like to acknowledge technical help from Katy McCortney, Emily Serbinowski, Lexie Trestan, the Cleveland Clinic Flow Cytometry Core and Dr. John Peterson from the Cleveland Clinic Imaging Core. We greatly appreciate the editorial assistance of Dr. Erin Mulkearns-Hubert. We would like to thank Amanda Mendelsohn from the Center for Medical Art and Photography at the Cleveland Clinic for the illustrations. Recombinant IL-2 was kindly provided by Dr. Marcela Diaz-Montero.

This work was supported by the National Institutes of Health R01NS109742 to J.D.L. and M.A.V., F32 CA243314-01 to D.B., and the Case Comprehensive Cancer Center Cancer Biology Training Award T32 CA059366 to D.B. This work was supported by National Institutes of Health grant R01NS102669 (C.H.), and by the Northwestern SPORE in Brain Cancer P50CA221747.

## Contributions

D.B., D.J.S., A.J.L., G.A.R., D.C.W., A.L., T.J.A. and B.O. carried out in vitro and in vivo animal experiments. Y.Z., C.P., C.H., D.V., A.M.K., T.H.H. and F.C. analyzed the sequencing data and performed bioinformatics analysis. D.B., M.M. and M.M.G. obtained and/or analyzed human data. M.D.S. and B.W.K. provided immunofluorescence analyses of human tissue. C.M.H. provided immunohistochemistry analyses of human tissue. M.A.V. C.M.H. B.W.K, A.M.K., T.H.H., M.S.A., F.C. and J.D.L. supervised the study. D.B. and J.D.L. conceptualized the study and wrote the manuscript. All authors read, revised, and approved the final manuscript.

