## Supplemental Figures for "Myeloid-derived suppressor cell subsets drive glioblastoma growth in a sex-specific manner"

**Supplementary Figure 1.** Increased mMDSC frequency in tumors associate with poor prognosis. **A,** Frequency of CD45$^{high}$ bone marrow-derived immune cells in resected tumors versus the contralateral hemisphere of sham-injected or 20,000 SB28-implanted mice on Day 14 and Day 21. Data shown as mean ± s.d. of n = 9-10/group *** $p<0.001$ as determined by two-way ANOVA. **B,** Percentage of mMDSCs (CD11b$^{+}$CD68$^{-}$Ly6C$^{+}$Ly6G$^{-}$I-A/I-E$^{-}$, CD11b$^{+}$CD68$^{-}$Ly6C$^{+}$Ly6G$^{-}$ or CD11b$^{+}$F4/80$^{-}$Ly6C$^{+}$Ly6G$^{-}$) and gMDSCs (CD11b$^{+}$CD68$^{-}$Ly6C$^{-}$Ly6G$^{+}$ or CD11b$^{+}$F4/80$^{-}$Ly6C$^{-}$Ly6G$^{+}$) in CD45$^{+}$ cells of the left hemisphere from n = 16 sham-injected and n = 26 GL261-bearing animals. Data shown for individual mice combined from three independent experiments and * $p<0.05$, ** $p<0.01$ as calculated by unpaired t-test. **C,** Percentage of mMDSCs in the left hemisphere 14 days post-SB28 implantation or sham injection. Data shown as mean ± s.d. from n = 5 sham-injected and n = 5 SB28-bearing mice per sex. ** $p<0.01$ as determined with unpaired t-test. **D,** Percentage of gMDSCs in the left hemisphere 14 days post-SB28 implantation or sham injection. Data shown as mean ± s.d. from n = 5 sham-injected and n = 5 SB28-bearing mice per sex. **E,** Ratio of mMDSCs to gMDSCs in the left hemisphere 14 days post-SB28 implantation or sham injection. Data shown as mean ± s.d. from n = 5 sham-injected and n = 5 SB28-bearing mice per sex. ** $p<0.05$, ** $p<0.01$ as determined with unpaired t-test. **F,** Percentage of mMDSCs and gMDSCs in the systemic circulation of n = 10 sham-injected and n = 10 SB28-bearing animals. Data shown for individual animals and ** $p<0.01$, *** $p<0.001$ as calculated by unpaired t-test. **G,** Kaplan-Meier curves depicting relative survival of male versus female mice orthotopically implanted with 25,000 GL261 cells. Data combined from three independent experiments with n = 15 females and n = 15 males. $p<0.05$ as determined by Gehan–Breslow–Wilcoxon test. **H,** Kaplan-Meier curves depicting relative survival of male versus female mice orthotopically implanted with 20,000 SB28 cells. Data representative of independent experiments with n = 5 females and n = 5 males. $p<0.05$ as determined by Gehan–Breslow–Wilcoxon test.

**Supplementary Figure 2.** Male to female bone marrow transplantation transfers the survival disadvantage observed in males. **A,** Kaplan-Meier curves depicting survival of female mice transplanted with bone marrow from male or female donors. Four weeks after the transplantation procedure mice were intracranially injected with 10,000 GL261 cells. Data shown from n = 11 male-to-female and n = 13 female-to-female bone marrow-transplanted mice. $p<0.05$ as determined by Gehan–Breslow–Wilcoxon test. **B,** Representative zebra plot demonstrating the efficacy of bone marrow transplantation at experimental endpoint. **C,** Ratio of mMDSCs to gMDSCs at experimental endpoint in blood and tumor of bone marrow-transplanted mice implanted with GL261. Data combined from n = 4 female-to-female and n = 4 male-to-female mice. $p<0.05$ as determined by unpaired t-test.

**Supplementary Figure 3.** The frequency of nonMDSC tumor-infiltrating immune populations is similar between males and females. **A-F,** Frequency of tumor-infiltrating immune populations 21 days post-GL261 implantation shown as mean ± s.d. **(A)** Percentage of macrophages (CD11b$^{+}$CD68$^{+}$ or CD11b$^{+}$F4/80$^{+}$) from n = 8-9 sham control mice and n = 13 tumor-bearing mice. Data shown from three independent experiments and * $p<0.05$ as determined by unpaired t-test. **(B)** Percentage of myeloid dendritic cells (CD11b$^{+}$CD11c$^{+}$) from n = 8-9 sham control mice and n = 13 tumor-bearing mice. Data shown from three independent experiments and *** $p<0.001$, * $p<0.05$ as determined by unpaired t-test. **(C)** Percentage of natural killer cells (CD3$^{-}$NK1.1$^{+}$) from n = 6-7 sham control mice and n = 9 tumor-bearing mice. Data shown from two independent experiments. **(D)** Percentage of total T cells (CD3$^{+}$NK1.1$^{-}$) from n = 6- sham control mice and n = 9 tumor-bearing mice. Data shown from two independent experiments. **(E)** Percentage of CD4$^{+}$ T cells (CD3$^{+}$CD4$^{+}$CD8$^{-}$) from n = 6-7 sham control mice and n = 9 tumor-bearing mice. Data shown from two independent experiments. **(F)** Percentage of CD8$^{+}$ T cells (CD3$^{+}$CD8$^{+}$CD4$^{-}$) from n = 6-7 sham control mice and n = 9 tumor-bearing mice. Data shown from two independent experiments. **G-L,** Frequency of tumor-infiltrating immune populations 14 days post-SB28 implantation shown as mean ± s.d. **(G)** Percentage of macrophages (CD11b$^{+}$CD68$^{+}$) from n = 5 mice/group. * $p<0.05$, *** $p<0.001$ as determined by unpaired t-test. **(H)** Percentage of myeloid dendritic cells (CD11b$^{+}$CD11c$^{+}$) from n = 5 mice/group. * $p<0.05$, ** $p<0.01$ as determined by unpaired t-test. **(I)** Percentage of natural killer cells (CD3$^{-}$NK1.1$^{+}$) from n = 2 sham control mice and n = 5 tumor-bearing mice. ** $p<0.01$ as determined by unpaired t-test. **(J)** Percentage of total T cells (CD3$^{+}$NK1.1$^{-}$) from n = 2 sham control mice and n = 5 tumor-bearing mice. * $p<0.05$ as determined by unpaired t-test. **(K)** Percentage of CD4$^{+}$ T cells (CD3$^{+}$CD4$^{+}$CD8$^{-}$) from n = 2 sham control mice and n = 5 tumor-bearing mice. * $p<0.05$ as determined by unpaired t-test. **(L)** Percentage of CD8$^{+}$

T cells ($CD3^+CD8^+CD4^-$) from n = 2 sham control mice and n = 5 tumor-bearing mice. * $p<0.05$ as determined by unpaired t-test.

**Supplementary Figure 4.** Increase in the systemic levels of immune cells is limited to gMDSCs. **A-F,** Frequency of circulating immune populations 21 days post-GL261 implantation shown as mean ± s.d. **(A)** Percentage of macrophages ($CD11b^+CD68^+$ or $CD11b^+F4/80^+$) from n = 9 sham control mice and n = 13 tumor-bearing mice. Data shown from three independent experiments. **(B)** Percentage of myeloid dendritic cells ($CD11b^+CD11c^+$) from n = 9 sham control mice and n = 13 tumor-bearing mice. Data shown from three independent experiments. **(C)** Percentage of natural killer cells ($CD3^-NK1.1^+$) from n = 7 sham control mice and n = 9 tumor-bearing mice. Data shown from two independent experiments. **(D)** Percentage of total T cells ($CD3^+NK1.1^-$) from n = 6-7 sham control mice and n = 9 tumor-bearing mice. Data shown from two independent experiments and * $p<0.05$, ** $p<0.01$ as determined by unpaired t-test. **(E)** Percentage of $CD4^+$ T cells ($CD3^+CD4^+CD8^-$) from n = 7 sham control mice and n = 9 tumor-bearing mice. Data shown from two independent experiments and * $p<0.05$, *** $p<0.001$ as determined by unpaired t-test. **(F)** Percentage of $CD8^+$ T cells ($CD3^+CD8^+CD4^-$) from n = 7 sham control mice and n = 9 tumor-bearing mice. Data shown from two independent experiments and * $p<0.05$, ** $p<0.01$ as determined by unpaired t-test.

**Supplementary Figure 5.** Inefficiency of mMDSC depletion is linked to active cell proliferation. **A,** Percentage of mMDSCs and gMDSCs from blood (top) and tumor (bottom) of tumor-bearing mice at experimental endpoint. Data shown as mean ± s.d. of n = 5-6 (2-3 male/3 female) for isotype-, n = 5-6 (3 male/2-3 female) for anti-Ly6G- and n = 6 (4 male/2 female) for anti-Ly6C-treated mice. *** $p<0.001$ as determined by unpaired Student's t-test. **B,** Fold difference in Ki-67 mean fluorescence intensity (MFI) of mMDSCs versus gMDSCs isolated from blood and tumor of n = 4 mice 21 days tumor implantation. Data shown as mean ± s.e.m. **C,** Comparison of Ki-67 intensity of mMDSCs isolated from n = 3 mice. Data shown as mean ± s.d. and statistical analysis performed with paired t-test.

**Supplementary Figure 6.** In vitro generated mMDSC and gMDSC have different gene expression profiles. **A,** Comparison of the polarization efficacy of mMDSCs and gMDSCs generated via co-culture approach or by cytokine stimulation. Data shown as mean ± s.d. from n = 3 females (left) and males (right). **B,** Volcano plots demonstrating the differential gene expression signature of mMDSC, gMDSC and nonMDSC fractions from n = 6 biological replicates. **C,** Number of differentially upregulated genes in mMDSCs (top) and gMDSCs (middle) in comparison to the other MDSC subset and control. Genes upregulated in both MDSC subsets (bottom) from n = 6 biological replicates. **D,** Top 10 differentially regulated pathways between mMDSCs versus gMDSCs using GeneOntology Enrichment Analysis. **E,** Comparison of the specificity of fludarabine to the chemotherapeutic agents used to target immunosuppression in GBM (Otvos et al., *Stem Cells*, 2016 & Alban et al., *JCI Insight*, 2019). **F,** Specificity of the multiple IL-1 pathway inhibitors predicted to target gMDSCs with network medicine analysis. **G,** Fold difference in IL-1β expression in MDSC subsets compared to nonMDSCs. Data shown as mean ± s.d. from n = 3 male versus female mice. $p<0.05$, ** $p<0.01$ as determined by unpaired t-test.

**Supplementary Figure 7.** MDSC gating strategy with flow cytometry and expression levels in the TCGA database. **A,** Gating strategy for the analysis of tumor-infiltrating mMDSCs and gMDSCs from patients with GBM. LOX-1 positivity of gMDSCs as determined based on expression levels in mMDSCs. **B,** OLR1 TPM expression values from the TCGA GBM dataset. Data shown as box and whiskers from n = 107 male and n = 59 female patients. Comparison is based on unpaired Student's t-test. **C,** IL-1β TPM expression values from the TCGA GBM dataset. Data shown as box and whiskers from n = 107 male and n = 59 female patients. Comparison is based on unpaired Student's t-test.

### Supplementary Figure 1

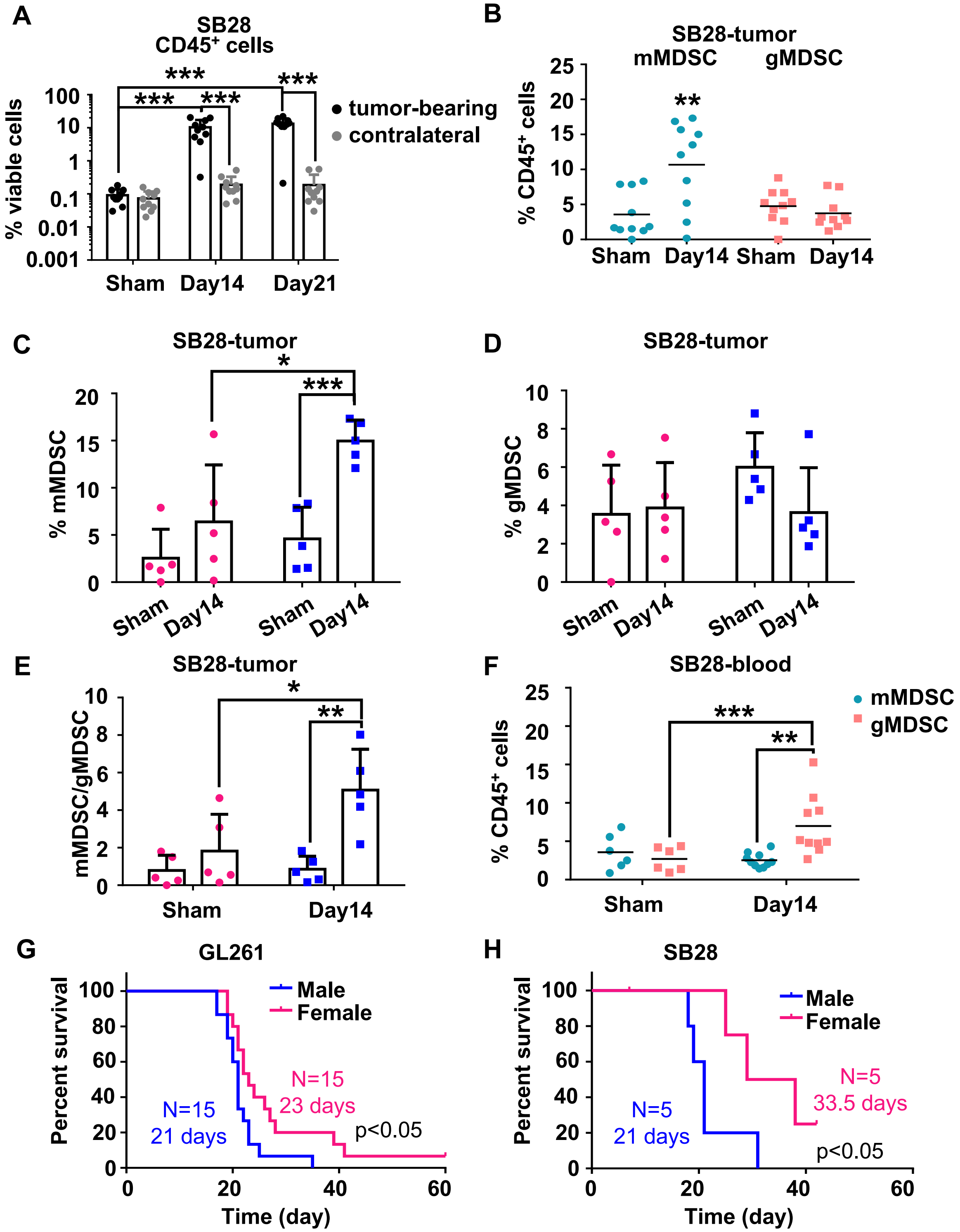

### Supplementary Figure 2

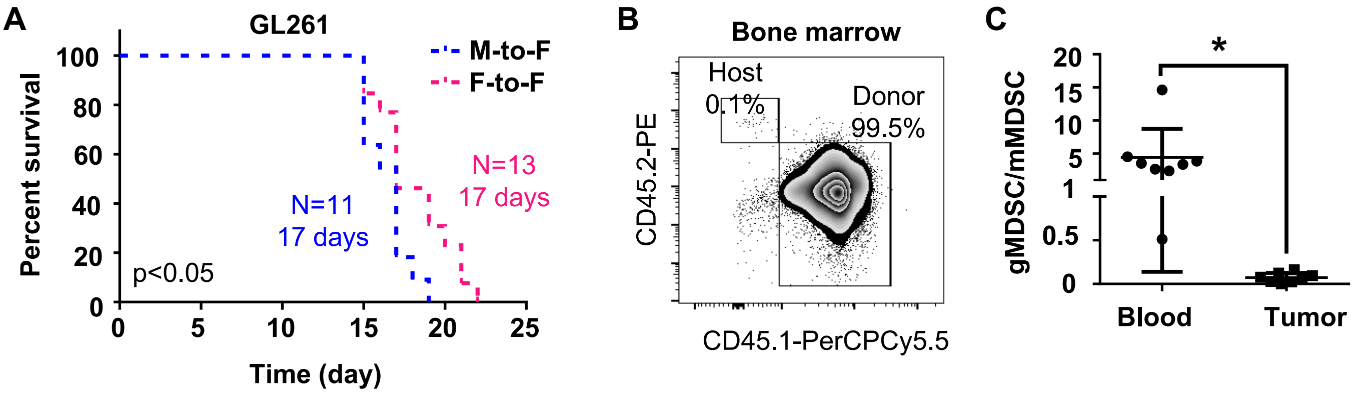

### Supplementary Figure 3

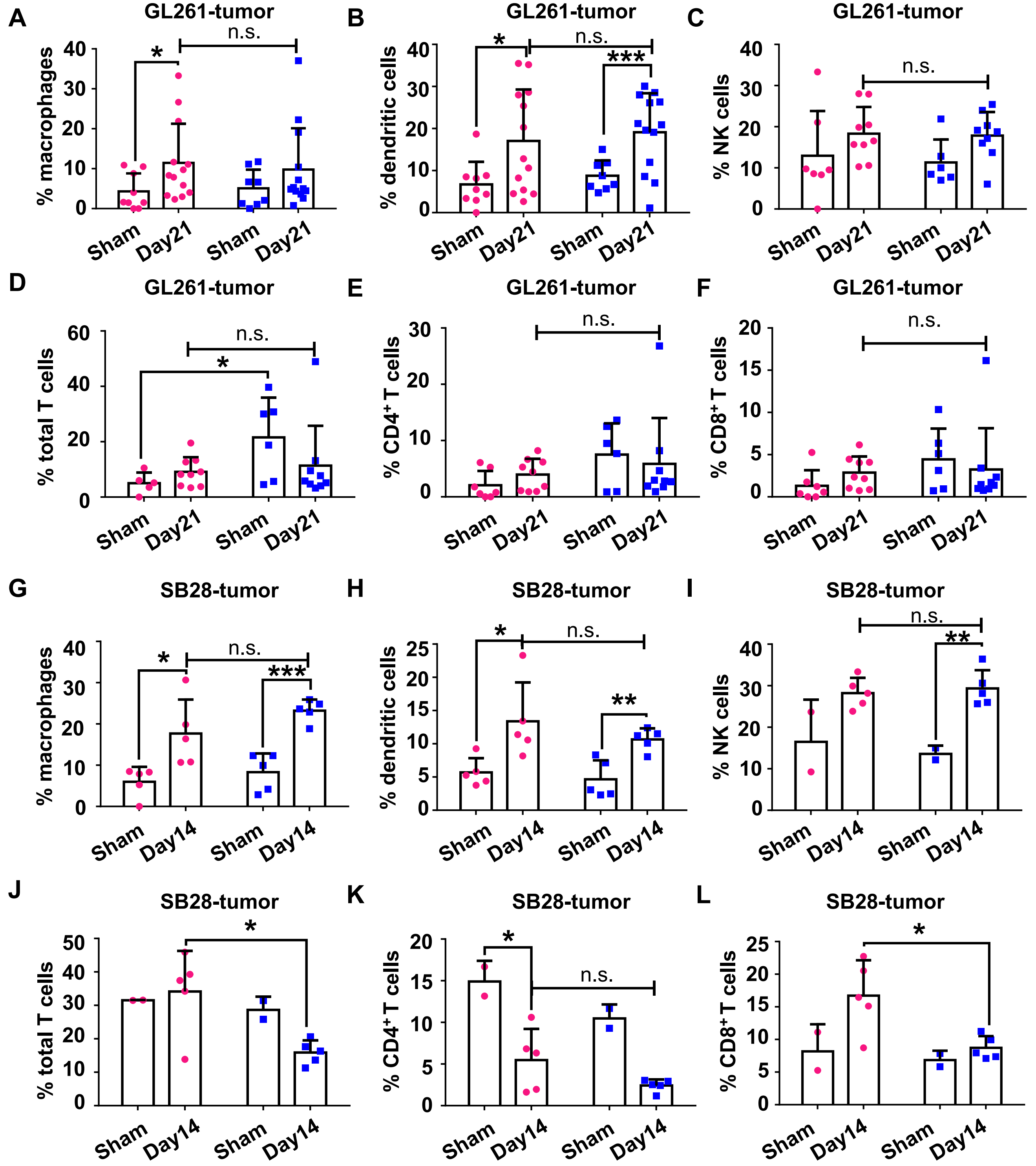

### Supplementary Figure 4

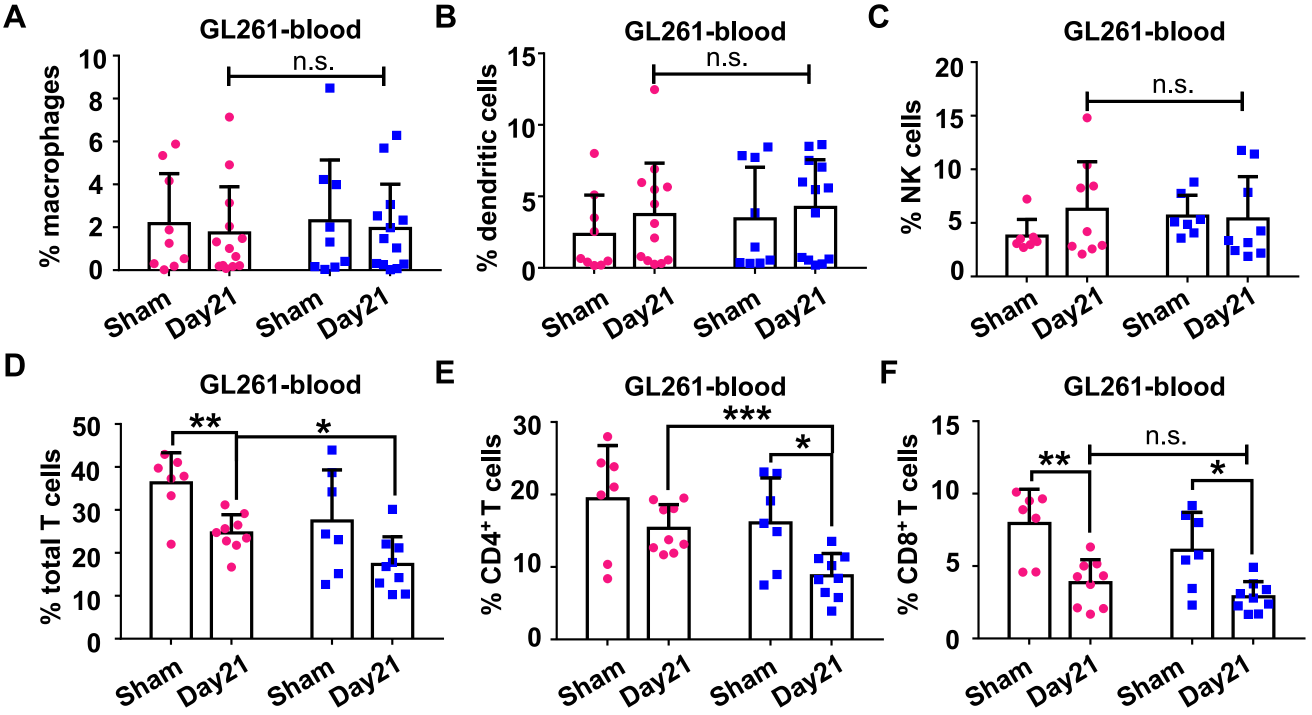

### Supplementary Figure 5

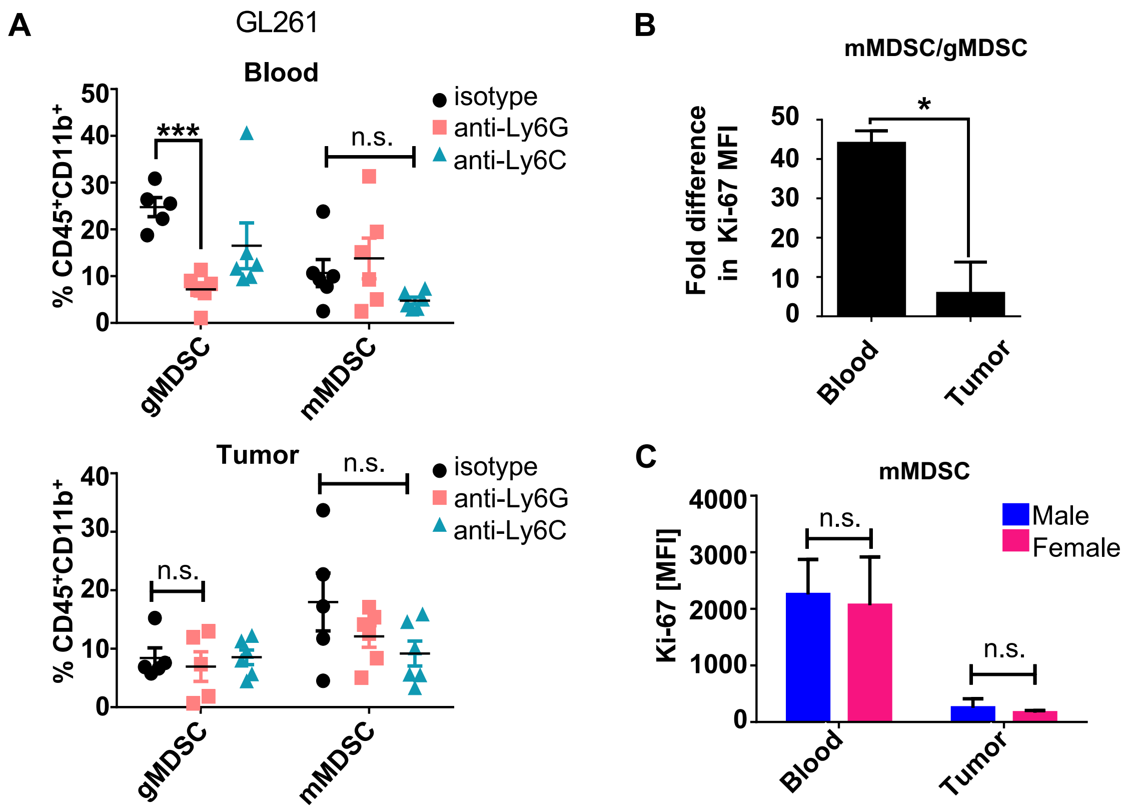

### Supplementary Figure 6

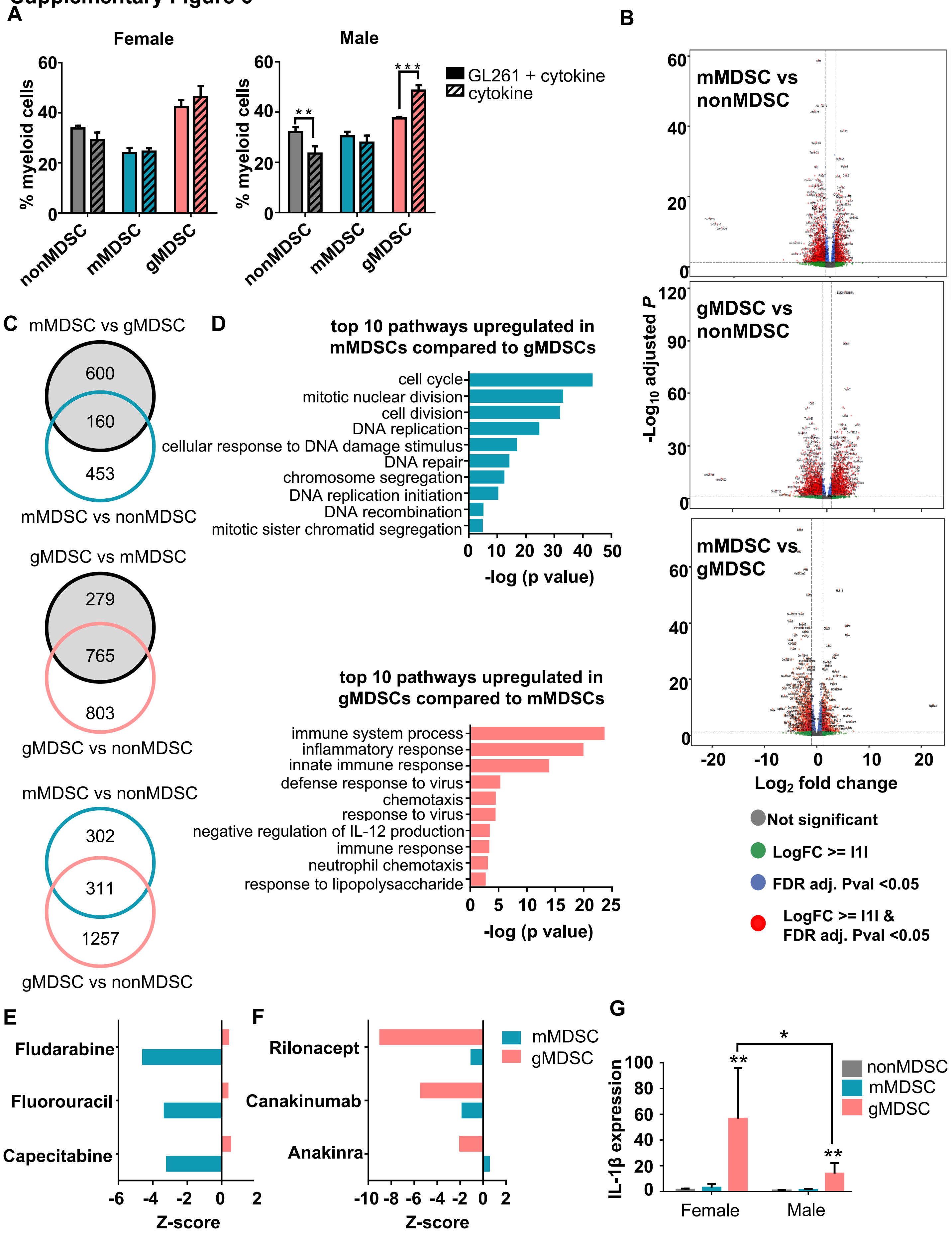

**Supplementary Figure 7**

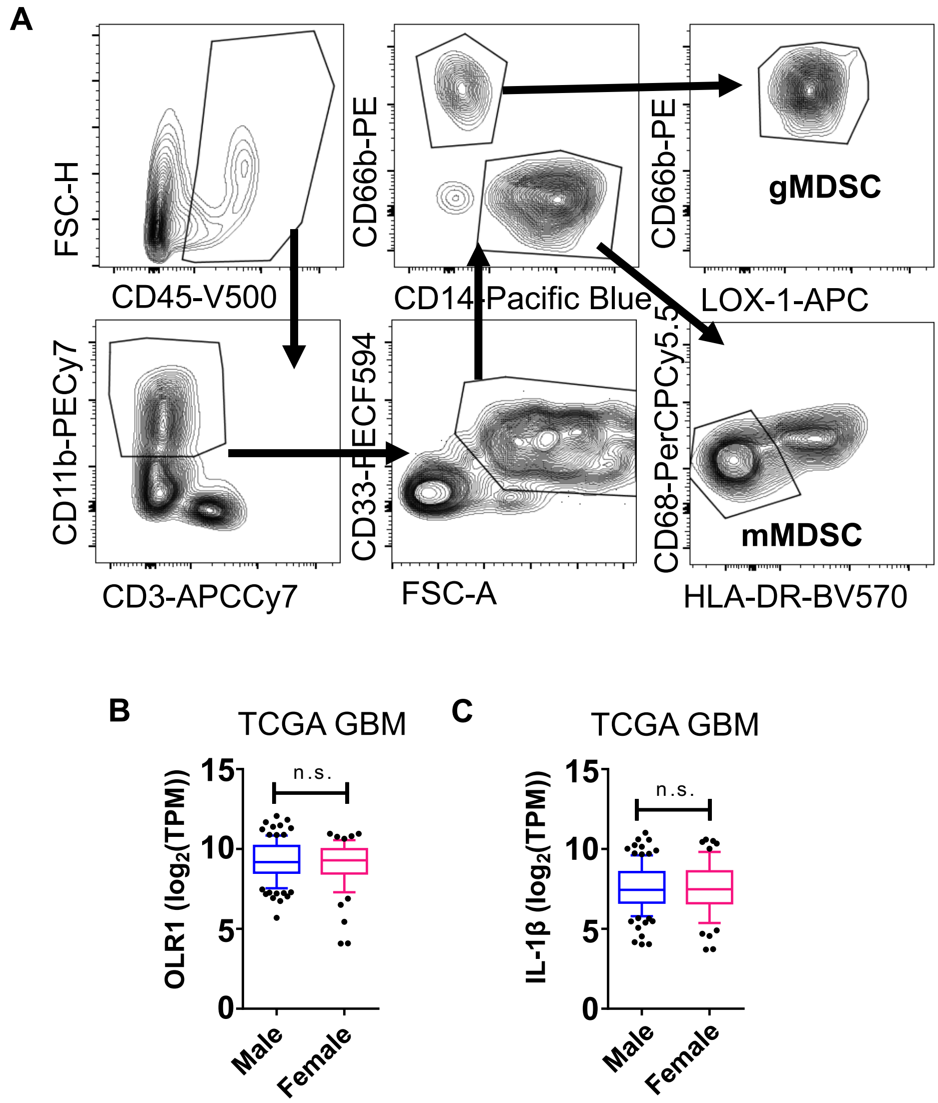
